# Onsager’s relations and Ecology

**DOI:** 10.1101/2023.11.28.569100

**Authors:** Jae S. Choi, Roger I.C. Hansell

## Abstract

There are two complementary approaches to thermodynamics: an empirical, phenomenological representation of macroscopic states and a model-based, statistical-mechanical representation of microscopic states. When only a few energy transformation steps are involved, macroscopic quantities such as free energy and entropy can be estimated unambiguously alongside well-characterized microscopic states. Both approaches have guided foundational ideas in ecology, yet natural ecosystems involve a vast number of coupled transformations across wide spatial, temporal, and organizational scales, leaving whole-system boundaries ill-defined and keeping microscopic and macroscopic representations from converging. This review clarifies the scope and limitations of information-theoretic order principles and extensive free-energy budgets in ecology, and synthesizes Onsager’s non-equilibrium reciprocal relations within a local, intensive control-volume framework (V^‡^), showing how they connect with the Le Chatelier-Braun Principle.

## 1 Introduction^1^

A coherent understanding of empirically observed *thermodynamic* states of matter was developed by Carnot (1824), Thomson (1849), and Clausius (1850). It was a monumental feat in that only a few variables of state (temperature, pressure, volume) were sufficient to describe the behavior of matter even if composed of an enormous number of atoms and molecules. Shortly thereafter, Boltzmann (1877) and Gibbs (1902) developed a statistical-mechanical description of the component parts that was complementary and related to the former by a proportionality factor, Boltzmann’s constant (Planck 1901).

All ecological processes are subject to these same thermodynamic laws.^2^ Even the most basic attempts to close matter and energy budgets necessarily invoke conservation of mass and energy (1^st^ law) alongside irreversible metabolic heat dissipation across system boundaries (2^nd^ law). There have been many attempts to understand this connection and draw inferences from it, motivated by the possibility of similarly and dramatically reducing the very large degrees of freedom required to describe ecological systems into a few variables of ecological “state”, and perhaps, even understanding the directionality of ecological dynamics.

Unfortunately, most ecological systems exhibit matter and energy cycling and boundary flows that occur at many space and time scales. This makes accounting for *entropy* very difficult or impossible (Meysman et al. 2010). As such, many resort to a model-based, information-theoretic representation of microscopic variability as an analogue to microscopic *disorder*. Further, due to the rather spectacular successes of Boltzmann (1877), many authors conflate the two concepts of macroscopic entropy and microscopic disorder. Indeed, it is only when both the microscopic and the macroscopic descriptions of a given system in focus can be shown to converge that one can begin to suggest that the system is reasonably understood. In molecular motion, certainly this connection has been demonstrated and so one says it is “understood”. But in ecological systems, both microscopic and macroscopic representations are far from convergent. It is, therefore, necessary to keep these two streams separate as they carry different perspectives, and one must remain cognizant of how much, or how little, they converge in order to gauge how well an ecological system is truly understood.

This paper reviews some of the key issues related to the application of thermodynamics in the field of ecology (Chapman et al. 2016; see also Fath et al. 2004; Chakrabarti et al. 2009; Meysman et al. 2010; Yen et al. 2014; Choi 2019). In particular, the discussion touches upon the conceptual scope and challenges of information-theoretic Order Principles; the macroscopic, extensive focus of free-energy gradient dissipation approaches and their associated limitations; and finally upon Onsager’s (1931a, 1931b) far-reaching contributions connecting the microscopic with the macroscopic, which have still not yet been sufficiently appreciated by ecologists, even after almost a century since their formulation. A companion paper (Choi 2026) develops the formal dynamical mapping between ecological community matrices and Onsager conductivity matrices.

## 2 Microscopic Order

Boltzmann (1877) connected the logarithm of the number of possible configurations of microscopic states in the phase space (statistical-mechanical) representation of a system to its macroscopic state variable, *entropy*, through a proportionality factor (Boltzmann’s constant *k*_B_). Boltzmann (1877) and subsequently Gibbs (1902) suggested that this increase in the number of microscopic states in phase space represents the amount of randomness or variability in potential microscopic states. The canonical example is an isolated container in which *n* molecules of a gas are initially confined by a partition to a small subvolume with a narrow range of momenta. Because work was required to compress and constrain the molecules into this restricted configuration, the accessible phase-space is small, the free energy available to do work is high, and the thermodynamic entropy sits at a low (reference) value. Once the internal partition is removed, molecular motion and collisions rapidly disperse positions across the full container and broaden the momentum distribution toward the Maxwell-Boltzmann equilibrium. Total internal energy is strictly conserved (1^st^ law), yet the number of accessible microscopic configurations in the 6*n*-dimensional phase space expands to a maximum (*order* → *disorder*) with a corresponding irreversible increase in macroscopic *entropy* and loss of available work.

Similarly, an internal combustion engine releases the chemical potential energy of fuel and converts it into mechanical work, dispersed exhaust gases, small particulates, and heat, generating thermodynamic *entropy* (Δ*S* ≥ *δQ/T*) and reducing the energy available to perform further work. From a microscopic perspective, considering the physical-chemical model of long-chain hydrocarbons and associated bond energies, the fuel transitions from a structured to a disaggregated state; that is, the number of accessible microscopic configurations in phase space has increased (*order* → *disorder*).

Schrödinger (1944) pointed out that biological systems seem to operate in a different manner. The example he used was DNA, which he described as a curious “aperiodic crystal” of extremely high information content, that is, with all the information required to create and maintain life. He noted that hereditary material manages to replicate itself with extreme stability as mutation rates are much lower than might be expected simply due to thermal replacement and degradation in chemical reactions of similarly large molecules (Drake et al. 1998). If DNA is represented heuristically as a phase space with dimensionality equal to the number of base pairs, with each replication from parent to offspring the structural sequence order remains conserved and the range of variability at each base pair is constrained to be within a narrow range relative to the parental generation. Thus, the accessible functional configurations remain narrow, stable, and robust, exemplifying an *order* → *order* process.

This local stability is maintained by chemical activation energies and covalent bond stability, proofreading enzyme systems and repair proteins, and Darwinian selection. Indeed, stable life cycles, developmental canalization, phenotypic plasticity, ordered metabolic pathways such as the tricarboxylic acid (Krebs) cycle, predator-prey interactions, host-parasite interactions, habitat-niche partitioning, animal migration patterns, and spatial species distributions all represent examples of *constrained* dynamical processes of *order* → *order*. These are the mainstay of biology and ecology: feedback cycles and their regulatory inter-relationships (Yodzis 1981, 1988). More generally, such open self-maintaining configurations are known as *dissipative structures* (Prigogine 1967; Glansdorff et al. 1971; Glansdorff et al. 1974; Nicolis et al. 1977); they describe locally stable, *order* → *order* systems that maintain their internal structure and low local entropy at the cost of continuous free-energy degradation and entropy export to their surrounding environment. When the accounting boundary is expanded to encompass both the biological structure and its surrounding environment, the overall thermodynamic effect is *(local order* ↔ *local order)* → *(global disorder)*, drawing high-grade free energy in and exporting entropy out. Physical and chemical analogues include turbulent energy cascades (Kolmogorov 1941), Rayleigh-Bénard convection cells, Belousov-Zhabotinsky oscillating chemical waves (Zaikin et al. 1970), and reaction-diffusion morphogenesis patterns (Turing 1952; Glansdorff et al. 1971; Nicolis et al. 1977). Living systems, in the act of growth, maintenance, and reproduction, tap into external matter and energy flows to sustain local order (Bertalanffy 1950; Zotin et al. 1967).

Schrödinger (1944) also considered the steps required for the origin and diversification of life, from simple prebiotic compounds to autocatalytic matter-energy cycles and eventually the evolution of template replication mechanisms involving RNA, DNA, proteins, and membrane compartmentalization in the presence of persistent free-energy gradients (*hypercycles* of Eigen et al. 1979; Wicken 1980). These transitions illustrate a third motif, *disorder* → *order*, which has been widely discussed across evolutionary and systems biology (Lotka 1922a, 1922b; Schrödinger 1944; Bertalanffy 1950; Eigen et al. 1979; Wicken 1980; Wiley et al. 1982; Schneider et al. 1994; Schneider et al. 2005). Over evolutionary time, living systems have accumulated hierarchical layers of structural and metabolic organization relative to primordial replicators. In other words, Darwinian selection acts in the short term to stabilize functional organization (*order* → *order*) and over evolutionary timescales to assemble novel organizational constraints (*disorder* → *order*) through the accumulation of survival and foraging strategies in a variable abiotic and biotic environment (Van Valen 1973, 1976; Bell 1982).

This apparent tension between local biological organization (*[dis]order* → *order*) and the 2^nd^ law expectation of increasing total *entropy* (*order* → *disorder*) has been proposed by several authors as a thermodynamic signature distinguishing living from non-living systems. Lovelock (1972) and Lovelock and Margulis (1974) used this logic to formulate the “Gaia hypothesis”, proposing that planetary-scale atmospheric homeostasis (a persistent non-equilibrium chemical steady state) serves as an indicator of life on a planet; a concept subsequently formalized in planetary-scale thermodynamic Earth-system models (Kleidon 2010). Analogous thermodynamic inferences have been developed for ecological succession: the directional patterns of colonization, biomass accumulation, and trophic network elaboration across landscapes over ecological timescales (Elton 1927; Whittaker 1953; Odum et al. 1955; Hutchinson 1959; Margalef 1963, 1968; MacArthur et al. 1966; E. P. Odum 1969; Gladyshev 1978; Matsuno 1978; Johnson 1981; Kay 1991; Vanriel et al. 1995; Washida 1995), as well as for ontogenetic and developmental trajectories (Zotin et al. 1967; Lurié et al. 1979; Briedis et al. 1984; Aoki 1989; Holdaway et al. 2010).

The coexistence of these three order principles has generated persistent confusion when frames of reference are left implicit. Resolving this ambiguity requires making explicit the focal representation (microscopic statistical configurations versus macroscopic energy and matter pools), the boundary constraints and reference states, and the spatial, temporal, and organizational scales of observation.

To illustrate how these three regimes coexist within a single physical cascade, consider a parcel of water flowing downhill along multiple paths and streams toward the equilibrium sink (Fig. 1, region A). A tagged volume of water in an elevated lake initially occupies a state of high gravitational potential energy, low macroscopic kinetic energy, and uniform temperature and pressure. As it flows downhill, gravitational potential energy is converted into macroscopic kinetic energy and progressively dissipated through viscous friction, acoustic noise, sediment erosion, percolation into groundwater, and evaporation, generating thermodynamic entropy. Eventually, the water reaches sea level (the equilibrium reference state for gravitational potential), where the initial potential energy has been fully dissipated into thermal molecular motion, generating and exporting thermodynamic entropy. Simultaneously, the initial spatial localization of the water parcel disperses across atmospheric vapor, river channels, groundwater aquifers, and the ocean (*order* → *disorder*).

**Fig. 1:**
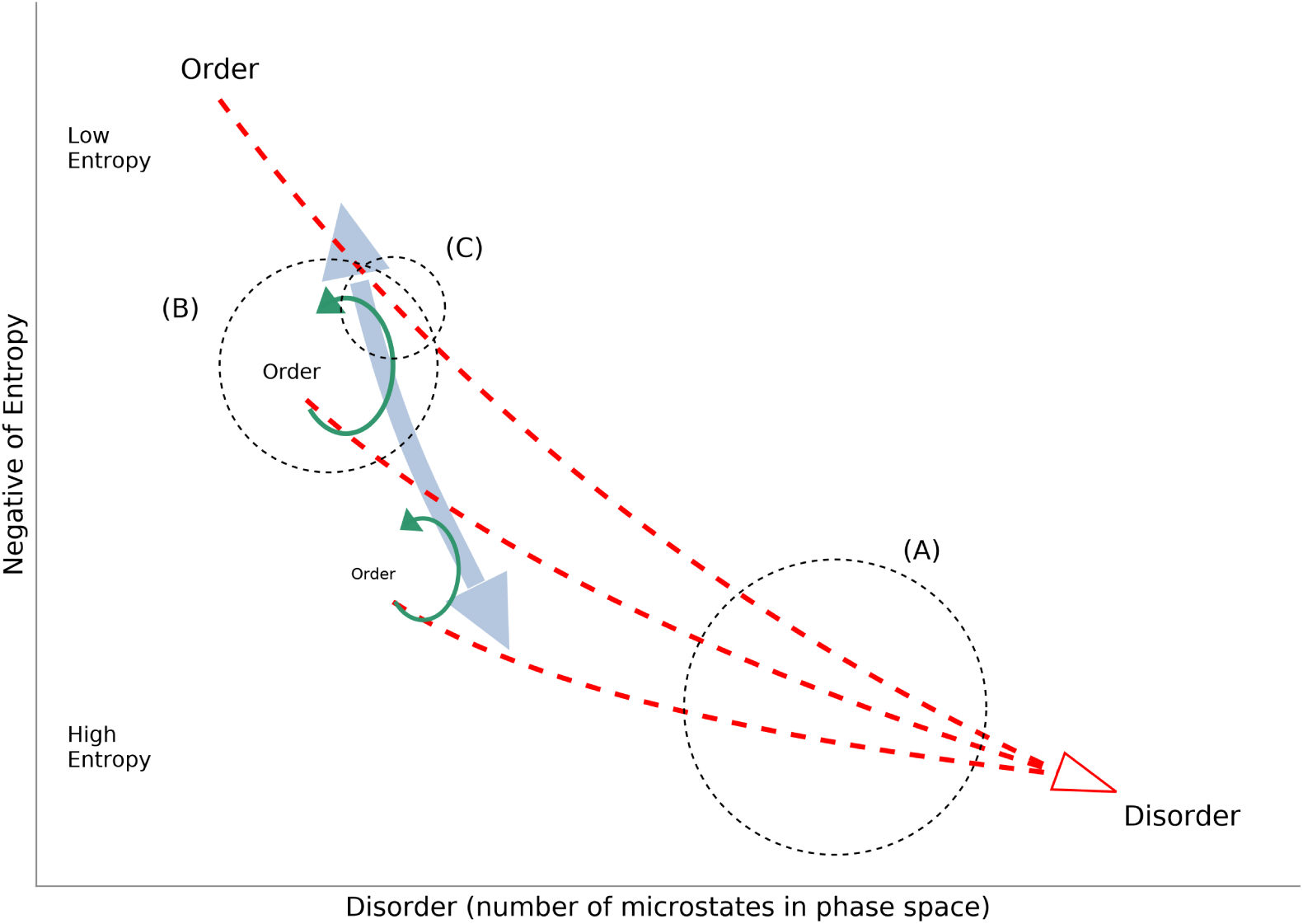
Heuristic 2D representation of the principles of order relating macroscopic negative entropy (Low Entropy to High Entropy, vertical axis) to microscopic disorder (number of microstates in phase space, horizontal axis). Red dashed trajectories represent irreversible free-energy degradation and entropy export from high Order toward equilibrium Disorder. Green recirculating arcs illustrate locally maintained dissipative structures across hierarchical tiers, linked by the shaded blue double-headed arrow. Dashed circles denote distinct frames of observation: (A) order → disorder along converging dissipative trajectories, (B) order → order within a stationary recirculating eddy, and (C) disorder → order across the branching transition from downward flow into structured cycling. Adapted from Choi (2019).

However, within specific spatial and temporal windows of observation, portions of the flow organize into persistent recirculating vortices, such as eddies and pools below rapids and falls. Within the local frame of these smaller space-time scales, such cycling structures maintain a stationary geometry and steady local order: *order* → *order* (Fig. 1, region B). Only a fraction of the water participates in the recirculating eddy at any instant while the bulk throughput continues downhill, exporting entropy and providing the continuous momentum gradient required to sustain the vortex against viscous dissipation. In biology and ecology, empirical research focuses primarily on characterizing the boundary conditions and internal reciprocal couplings (formalized below via Onsager’s coefficients *L_ij_*) that sustain analogous recirculating matter and energy structures across biochemical, organismal, and ecosystem tiers.

In ecology, there is an implicit understanding that these matter and energy flows are borrowed by biota, redirecting a small fraction of energy and matter into living structures through growth and reproduction before eventual metabolic dissipation. Living cycling structures depend upon and accelerate boundary energy degradation by exporting entropy via respiration, evapotranspiration, and chemical degradation. Lindeman (1942) was among the first to quantify trophic transfer losses; his measurements and subsequent syntheses by H.T. Odum (1956), E.P. Odum (1969), and Mann (1972) established that ecological transfer efficiencies generally range from 1 to 10%. Consequently, dissipative respiratory and boundary losses dominate energy transformations in ecological networks. Satellite-based radiation budgets similarly indicate that in terrestrial plant canopies, >99% of absorbed solar energy is dissipated through evapotranspiration and respiration (Schneider et al. 1994; Schneider et al. 2005), processes that perform essential physiological work by drawing water and dissolved nutrients from the soil while exporting entropy to the atmosphere. Only a small fraction of gross energy input is sequestered as standing biomass, yet over geological timescales this persistent biological flux substantially alters planetary conditions (Lovelock et al. 1974).

Finally, if the frame shifts to the transition from downward flow to recirculating structures (Fig. 1, region C), structured vortices emerge once inertial driving exceeds viscous damping past a critical threshold, inducing a bifurcation to a new dynamical attractor. Analogous regime shifts occur in ecological and evolutionary dynamics when external free-energy gradients or internal feedback loops cross stability thresholds due to environmental forcing, species invasions, or trophic cascades.

A rigorous evaluation of microscopic *order* and *disorder* requires an explicit dynamical and statistical *model* of the system. Because information-theoretic measures depend on the chosen state variables, coarse-graining categories, and parameterization, purely formal entropy indices applied to ecological models such as Lotka-Volterra interaction models (Michaelian 2005; Chakrabarti et al. 2009) or aggregate input-output flow networks (Patten 1978) yield model-dependent descriptors rather than absolute thermodynamic state functions. Historically, it was precisely the quantitative discrepancy between classical statistical models of molecular and radiation states and empirical calorimetry that revealed missing physical degrees of freedom, motivating quantum statistical mechanics (Planck 1901; Schrödinger 1944; Jaynes 1957). In ecology, information-theoretic indices have been applied to trophic flow topologies (Ulanowicz et al. 1987; Ulanowicz et al. 2006), genome size and coding complexity (Jørgensen et al. 1995), phylogenetic branching (Wicken 1980; Wiley et al. 1982), and maximum-entropy species distribution modeling (Elith et al. 2011). While these statistical tools quantify network diversity, constraint, or occurrence probability, connecting them to physical thermodynamics requires mapping their coarse-grained states to physical energy and matter pools with explicit microstate multiplicities and transition energetics.

## 3 Macroscopic Entropy

Given the lack of satisfactory microscopic model representations of ecosystem phenomena, many have attempted instead to describe ecosystem processes in terms of macroscopic state variables. Such a whole system-level (*extensive*) approach is perhaps best exemplified by the engineering tradition which traces its origin directly to the founding work of Carnot (1824). The whole-system free energy available to do work is the state variable of interest, carefully controlled and measured. Engineering applications usually focus upon systems whose boundaries can be defined explicitly (for example, a machine or industrial production process) and revolve around well-defined forms of energy and matter transformation with an explicitly defined reference or ground state. These conditions permit repeated measurements of free energy and entropy, and transfer efficiencies can be calculated to permit cost-benefit or efficiency analysis of the whole system.

Following this macroscopic tradition, Lindeman (1942), E.P. Odum (1969), Kay (1991), Aoki (1993, 1995), Schneider and Kay (1994), Müller (1998), Choi et al. (1999), Schneider and Sagan (2005), and Jørgensen (2012) analyzed *extensive* free-energy gradients and their degradation across whole ecosystems. Aoki (1993, 1995) estimated whole-lake entropy production from radiation and heat budgets; Choi et al. (1999) examined dissipative signatures in plankton size spectra; and Schneider and Kay (1994) used satellite measurements of evapotranspiration and surface temperature as proxies for ecosystem energy degradation.

These extensive accounting approaches, however, suffer a difficulty similar to those faced in defining any quantity with many mass-energy transformation processes occurring at large ranges of spatial, temporal, and organizational scales of open, non-equilibrium systems with ill-defined boundaries and reference states, often arranged in cycles. Further, that which is a resource for one organism can be irrelevant for another or even a toxin. These complex relationships between extensive variables of state and system stability suggest that a naive application of extensive thermodynamics is problematic (Meysman et al. 2010).

To overcome or at least acknowledge some of these challenges, Patten (1978) attempted to unravel energy transformation cycles as a first-order linear expansion of input-output relations. Similarly, H.T. Odum (1996) attempted to trace all energy transformation steps starting from solar radiation to the component of interest, expressing it in terms of solar “emergy” (emergent energy) units. Generally, the conclusion is that macroscopic entropy production increases monotonically in accordance with the 2^nd^ law, and biological complexity might be associated with this throughput, but aggregate black-box accounting abstracts away the internal feedback mechanisms, regulatory responses, or direction of perturbation recovery that are also of interest. Resolving how internal ecological interactions respond to perturbations requires moving from aggregate whole-system accounting to the linear and local non-equilibrium formalism of Onsager (1931a, 1931b), Prigogine (1955), and de Groot and Mazur (1962).

## 4 Onsager’s Relations

Onsager’s (1931a, 1931b; see also 1953; 1997; 2014) remarkable contributions tied macroscopic entropy with microscopic fluctuation statistics for any generalized thermodynamic system. Here the discussion repeats some of this foundational development for the sake of clarity; and then focuses upon the fascinating relationship between Onsager’s relations (*Conjugate effects*) and the well-known Le Chatelier-Braun Principle (Bertalanffy 1950; Gündüz et al. 2014).

Onsager’s relations are a generalization of well-known empirical transport phenomena such as Fick’s law of diffusion, Fourier’s law of thermal conduction, and Ohm’s law of electrical current, expressed in terms of entropy production. More generally, in any thermodynamic system, spatial gradients of intensive variables (such as temperature, chemical potential, or concentration) act as thermodynamic forces *X_j_* that drive conjugate fluxes *J_i_*of extensive matter or energy. Near a stationary reference state where forces are small, fluxes can be expressed as linear functions of the forces:

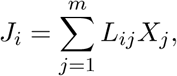

where *L_ii_* represents the direct conductivity of force *X_i_* on its own flux *J_i_*, and *L_ij_* (*i* ≠ *j*) represents cross-coupling between distinct physical processes.

With the assumption of microscopic reversibility of equilibrium fluctuations, Onsager proved that the matrix of phenomenological conductivities is symmetric:

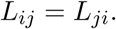

When irreversible transport processes are connected reciprocally in this fashion, they are designated *conjugate processes* exhibiting *Conjugate effects*. Confirmation of the reciprocal nature of these relationships in physical energy transformations has been verified experimentally across many systems (Miller 1960). A well-known, concrete physical example is the Seebeck effect where electrons flow in the presence of a temperature gradient, serving as the basis for thermocouples; and the subsequent discovery of the reciprocal relation, the flow of heat with the application of electricity, known as the Peltier effect, which serves as the basis for thermoelectric heat pumps and thermistors. For thermal-electromagnetic coupling, the relations are known as the Nernst-Ettingshausen effects. The same reciprocal couplings have been explored in chemical reactions and cell membrane dynamics (Kedem et al. 1958; Katchalsky et al. 1965; Oster et al. 1971; Mikulecky 1985).

The internal entropy production rate *σ* within an open system is non-negative and equals the sum of products of conjugate fluxes and forces (equivalently, a quadratic form in the forces):

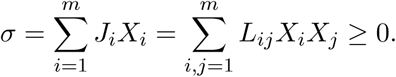

As a consequence, the symmetric conductivity matrix **L** must be positive definite, which also means all self-conductivities must be positive (*L_ii_ >* 0) and the cross-coefficients are bounded (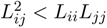) (Groot et al. 1962). In the linear near-equilibrium regime under fixed boundary forces, the stationary relaxation of unconstrained internal forces minimizes internal entropy production (Prigogine’s minimum entropy production theorem); and when forces are fixed, the realized fluxes extremize Onsager’s least-dissipation functional (Onsager 1931a, 1931b; Onsager et al. 1953; Shiner et al. 1994; Martyushev et al. 2006; Verhás 2014).^3^

When a system is subjected to an external perturbation *δX_i_* along a focal coordinate *i*, stabilization operates through two coupled responses: intrinsic primary self-damping (*L_ii_ >* 0) immediately opposes and dissipates displacement along the focal coordinate, while coupled secondary pathways (indirect pathways, sensu Patten 1978) adjust to absorb and redistribute stress across the network. This network cross-coupling moderates the effective conductance of the primary pathway via the matrix Schur complement (where the effective network-moderated conductivity satisfies 0 *< L_ii,_*_eff_ ≤ *L_ii_*), in alignment with the Le Chatelier-Braun principle in coupled transport networks (Prigogine et al. 1954; Bertalanffy 1950; Gündüz et al. 2014; see Choi 2026, for more details).^4^ Here, “primary” and “secondary” are relational designations rather than fixed species labels: every node acts as the primary dissipative channel for disturbances entering directly at its own coordinate, while simultaneously serving as a secondary moderating buffer for disturbances entering elsewhere in the network.

## 5 Conjugate Effects in Ecological Systems

Onsager’s relations formalize the Le Chatelier-Braun principle (*Conjugate effects*) within linearized non-equilibrium thermodynamics. In ecology, interactions such as mutualism (+*/*+), parasitism (+*/*−), and predation (+*/*−) are conventionally classified by the signs of the demographic community matrix **A**(Yodzis 1981, 1988; Michaelian 2005), which measures population growth-rate sensitivities (*∂Ḃ_i_/∂B_j_*). Those demographic signs are generally asymmetric (*A_ij_* ≠ *A_ji_*); and so, when they were asserted to be equivalent to Onsager conductivities (**A** ≡ **L**), researchers concluded that ecosystems violate Onsager’s relations and disregarded their utility. In reality, demographic rates conflate underlying physical transport conductivities with directional trophic cycling and heterogeneous standing biomass storage (the formal dynamical mapping is developed in Choi 2026).

However, in the statistical ensemble of ecosystem interactions, and not a single feeding event, ecologists understand that matter and energy exchanges occur recurrently around a stationary state. That is, they cycle, such that a local specific entropy production rate can be defined (at some spatial and/or temporal scale), *σ* = **X**^T^**LX** ≥ 0. From this perspective, ecological reciprocity is expressed through shared dissipative (metabolic) coupling in *σ* and network-moderated conductivity (0 *< L_ii,_*_eff_ ≤ *L_ii_*) rather than symmetric demographic coefficients in **A**. In tightly coupled metabolic symbioses, repeated bidirectional substrate and energy exchanges mean a shared dissipative flux-force relationship **L**, so perturbing the metabolic gradient of one partner induces a reciprocal counter-flux in the other.

In host-parasite and predator-prey systems, the asymmetric +*/*− demographic signs (in **A**) reflect directional trophic transfer and heterogeneous standing biomass storage, yet repeated interactions still generate stabilizing conjugate feedbacks in **L** and *σ*. Over ecological and evolutionary timescales, predation regularizes prey fluctuations and prevents boom-and-bust resource exhaustion. A classic example is white-tailed deer introductions to predator-free islands collapsing after depleting forage, whereas when predators exist, they can coexist (Russell et al. 2001). In host-parasite co-evolution (Van Valen 1973) and controlled immune stimulation (Strachan 1989, 2000; Rook 2009; Smith et al. 2012), analogous compensatory stabilization at multi-generational scales is evident.

At broader biogeochemical scales, the Gaia hypothesis (Lovelock 1972; Lovelock et al. 1974) and Daisyworld models (Watson et al. 1983) illustrate reciprocal feedbacks between organismal metabolism and planetary climate, atmospheric composition, and albedo. Similarly, statistical correlations among respiration, biomass production, photosynthesis, and evapotranspiration (Kleiber 1947; Bertalanffy 1950), benthic-pelagic coupling in lakes (Regier et al. 1996), and the marine biological carbon pump (Eppley et al. 1979) reflect mechanically coupled mass-energy fluxes whose causal reciprocity is testable: perturbing one driving gradient must elicit a measurable counter-response in its conjugate flux rather than mere statistical correlation/regression to the mean.

Even where compartments might not seem directly coupled, the presence of recurrent, indirect network pathways generates *effective* conjugate moderation via the Schur complement. For example, Halbach (1973) observed that rotifers at elevated temperatures mature at smaller body sizes with higher population oscillation frequencies. Under an effective thermodynamic coupling between thermal dissipation and size-structured biomass, holding temperature fixed while shifting mean body size predicts compensatory changes in numerical density so that total community heat dissipation adjusts toward the local balance— providing a falsifiable mechanistic prediction rather than a “just-so” descriptive correlation (Peters 1991). Conversely, strong positive feedbacks or sustained overexploitation can rupture established conjugate couplings, as seen in the collapse of the Northwest Atlantic cod stock and its transition to an alternative community regime (Choi et al. 2004; Choi 2022) (*order through fluctuation*, sensu Prigogine et al. 1972; Babloyantz 1986). Ecologists have long studied these compensatory feedbacks empirically; Onsager’s transport formalism situates them within a unified physical framework grounded in symmetric dissipative reciprocity (**L**) and specific entropy production (*σ*).

## 6 Local, Intensive Description

To apply irreversible thermodynamics to open, spatially heterogeneous ecosystems, Prigogine (1955) and de Groot and Mazur (1962) formulated Onsager’s relations as a *local, intensive* description within an arbitrary control volume element *V^‡^* (indicated by the superscript *^‡^*). To emphasize this nuance, the qualification (*local, intensive*) merits repetition. Its importance has often been overlooked in the literature, where system volume is assumed constant, conflating intensive with extensive expectations (frequently encountered because in most physical applications system volume does not change, so intensive and extensive variables are simply proportional). In living systems, macroscopic (*extensive*) biovolume fluctuations of many orders of magnitude make naive applications of whole-system Onsager relations problematic. A constant local spatial control volume element *V ^‡^*, within which living biomass and biovolume fluctuate, is however defensible, establishing the validity of the Onsager relations.^5^

Within a local volume element, the change in subsystem entropy *dS^‡^*over time *dt* is the sum of internal entropy production *d_i_S^‡^* ≥ 0 and boundary entropy exchange *d_e_S^‡^*:

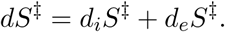

Dividing by control volume *V^‡^* defines the local specific entropy production rate *σ^‡^*:

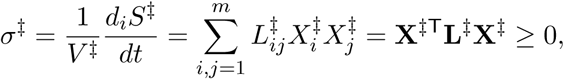

where 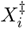 are local intensive thermodynamic forces and 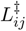 are local symmetric phenomenological conductivities (**L***^‡^* = **L***^‡^*^T^ ≻ 0).

This addition of an entropy transport process permits internal subsystem entropy to behave in ways that seem locally counter to the 2^nd^ law, though of course total entropy increases in accordance with physical law (Fig. 1). In the linear near-equilibrium regime, expectations of local Conjugate effects coincide with a local least specific dissipation of energy principle (Choi et al. 1999; Choi et al. 2001); extensions to strongly driven trophic networks are developed in the companion paper (Choi 2026). Here, the qualification “specific” emphasizes an expectation of local intensive variables. Operationally, expressing ecosystem respiration *R* and standing biomass *B* on a common areal control-column basis (m^−2^, crudely approximated by depth-integrating across the active water column in pelagic systems to match terrestrial and benthic cross-sections) provides the local control-volume (*V*^‡^) representation of *σ*^‡^ and storage capacitance, eliminating extensive domain-size artifacts while keeping kinetic turnover rates *R* = *R/B* (yr^−1^) invariant across areal and volumetric normalizations (Fig. 2).

**Fig. 2:**
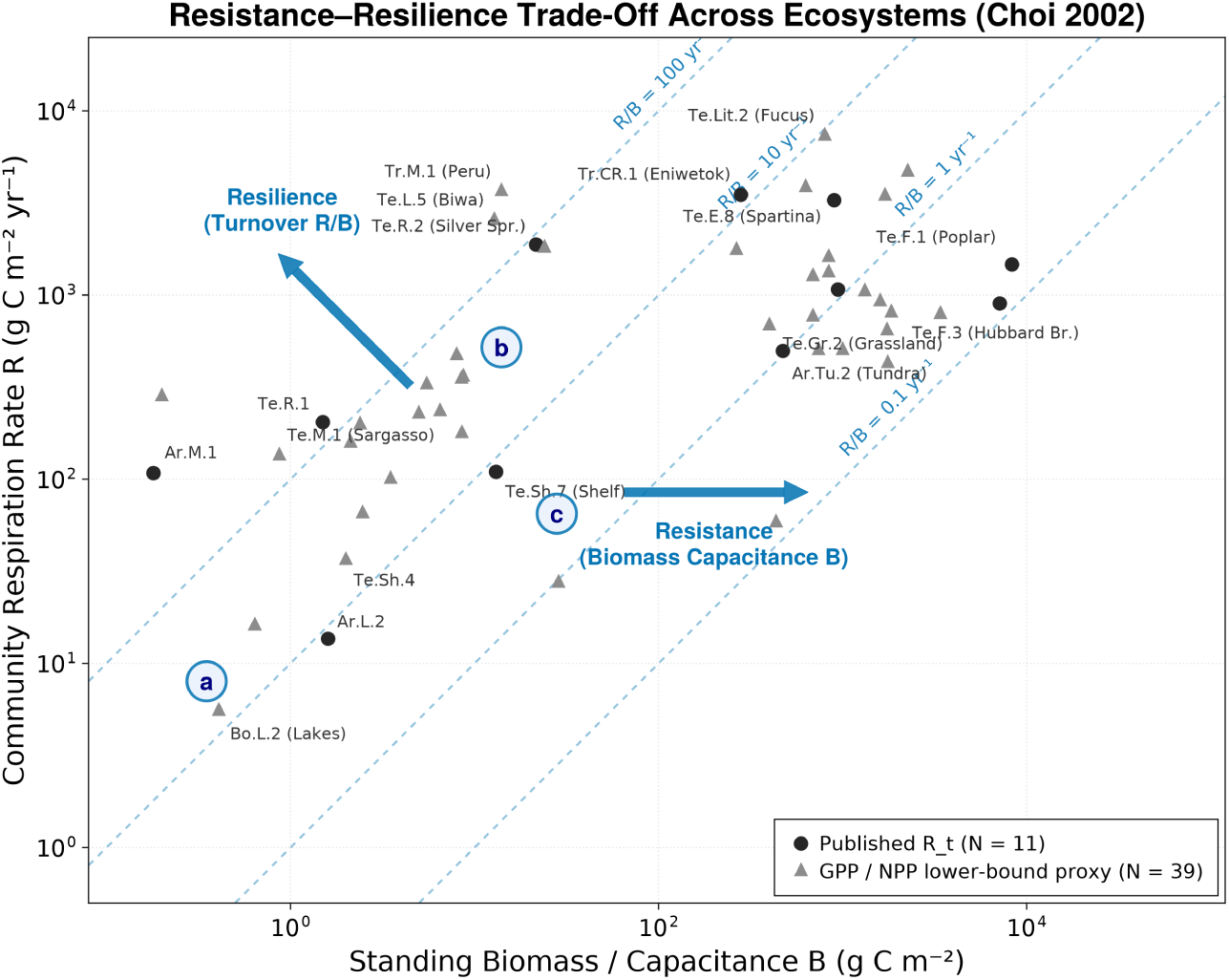
Resistance–Resilience trade-off between community respiration rate R (a proxy for entropy production) and standing biomass capacitance B across 50 ecosystems compiled in Choi (2002), comprising published total respiration R_t_ observations (black circles, N = 11) and GPP/NPP lower-bound metabolic proxies (gray triangles, N = 39). Parallel 45^◦^ dashed isoclines mark constant Resilience (turnover rate R/B ∈ {0.1, 1, 10, 100} yr^−1^). Blue vectors indicate Resilience (perpendicular to the constant-R/B isoclines) and Resistance (biomass capacitance B, parallel to the horizontal B-axis). Regions (a) and (b) illustrate dilute high-turnover versus high-biomass operating regimes, while local deviations such as (c) represent displaced configurations expected to relax toward the cross-biome trade-off band (see Choi 2026, for the expanded multi-biome synthesis, nonlinear crossover fits, and thermal normalization). Climatic regions: Te=temperate; Tr=tropical; Ar=arctic; Bo=boreal. Ecological types: E=estuary; L=lake; Lit=littoral; M=marine; Sh=continental shelf; CR=coral reef; R=river; Tu=tundra; Gr=grassland; F=forest.

For general theorists, this distinction is often considered minor because they integrate across volume elements and expect aggregate behavior to be concordant. This holds when systems under study (such as cell membranes or atmospheric columns) maintain near-constant total volume. Ignored, of course, is the variability of biovolume, which has its own dynamic and structural properties in multicellular and ecological systems; consequently, macroscopic and microscopic thermodynamic processes may not be concordant (Fig. 2). Internal structural transients and lags that appear negligible from a macroscopic perspective may be decisive across the space-time scales of organisms and human observers.

In the intensive approach, there is an explicit expectation of local Conjugate effects, that is, local steady states or local homeostasis (Fig. 2). At an organismal level, this expectation of homeostasis is not considered surprising by medical and physiological sciences as it is a necessary constraint for the continuity of life. However, at population, community, evolutionary, or planetary scales, arguments relating to homeostasis have frequently been met with skepticism and pejoratively labeled as “vitalism”, “Lamarckism”, “Panglossianism”, or “teleological” (Tansley 1935; Gould et al. 1979; Mayr 1991; Grimm et al. 1997). The importance of the non-equilibrium thermodynamic perspective is that it provides a local directionality of change that is consistent from organismal to subcellular to Earth-system scales, without resorting to teleology: an expectation grounded simply in the directionality of time and the dissipation of free-energy gradients along kinetic pathways.

Prigogine (1955) and Prigogine et al. (1972) coined the term “dissipative structures” to describe open, non-equilibrium systems that maintain internal spatial and temporal order by continuously importing high-grade free energy, performing metabolic work, and exporting degraded entropy to their surroundings (−*d_e_S^‡^* ≥ *d_i_S^‡^* ≥ 0). In ecosystems, these structures are organized hierarchically across scales, from subcellular organelle cycles and whole-organism physiology to population dynamics, food-web topologies, and biogeochemical cycles (O’Neill et al. 1986; Choi et al. 2001). Although the flow of energy across trophic size spectra has frequently been likened to fluid turbulence (Kolmogorov 1941), the biological cascade possesses a different physical architecture. In fluid mechanics, kinetic energy cascades strictly unidirectionally from large eddies down through an inertial subrange to the viscous microscale, where thermal dissipation exclusively occurs. In living communities, by contrast, metabolic dissipation occurs across all trophic levels and body-size classes through cellular respiration, while matter and energy circulate multidirectionally across the size spectrum via predation, detrital recycling, and parasitism.

Because mass-specific metabolic rate scales inversely with organismal body mass according to Kleiber’s law (Kleiber 1947; Bertalanffy 1950; Peters 1983; Choi et al. 1999), these conjugate compensations operate across the allometric size spectrum. Large-bodied organisms accumulate substantial standing biomass and provide high storage capacitance (conferring resistance against environmental fluctuations), whereas small-bodied organisms exhibit rapid mass-specific turnover (conferring kinetic resilience and high recovery speed). Dimensionally, [*R/B*] = T*^−^*^1^ defines an inverse relaxation timescale (R = *R/B*) that decreases as system size, mean organismal mass, and geometric packing increase (Fig. 2). Across the full biomass spectrum, this scale dependence implies a transition from kinetic-limited turnover in dilute planktonic systems (*R* a *B*^1^ along high-*R/B* isoclines) through an allometric regime (*R* a *B*^1^*^/^*^4–1^*^/^*^3^ as larger-bodied structural biomass accumulates) toward external solar and nutrient boundary-flux saturation in dense forests (*R* → *R*_max_), a crossover illustrated qualitatively across biomes (Fig. 2) and evaluated quantitatively across an expanded multi-biome synthesis (see Choi 2026). Perturbing the size structure or abundance of an apex predator guild necessarily induces reciprocal shifts in the size distribution and turnover rates of prey and competitors. When total community biomass is constrained by environmental carrying capacity, total metabolic throughput must adjust; conversely, when energetic input is bounded by boundary flux, standing biomass and species composition reorganize. If both biomass and energy inputs are clamped, organisms must adapt through altered metabolic efficiencies, life-history shifts, or phenotypic plasticity; in this way, physical conservation laws constrain size-spectrum dynamics and life-history evolution.

Beyond the linear near-equilibrium regime, extending thermodynamic stability analysis into non-linear dynamics requires accounting for complex state-space topologies (Nicolis et al. 1989). Far from equilibrium, universal variational extrema no longer hold unconditionally; non-linear ecological networks exhibit multiple stability domains, semi-stable attractors, and threshold bifurcations (Shiner et al. 1994; Byers et al. 1996). Under strong external gradients, macroscopic organization emerges via Prigogine’s *order through fluctuation* mechanism (Prigogine et al. 1972; Nicolis et al. 1977; Nicolis et al. 1989): when non-linear feedbacks drive a thermodynamic branch past a critical instability threshold, spontaneous internal fluctuations are amplified rather than damped, carrying the system into a new, spatially or temporally structured dissipative regime without violating microscopic fluctuation relations (Crooks 1999).

In ecology, early attempts to capture this directional tendency culminated in the Maximum Power Principle (Lotka 1922a, 1922b; Odum et al. 1955), which hypothesized that natural selection maximizes useful energy throughput. Odum and Pinkerton (1955) noted the quadratic form of power output across energy transformations, arguing that systems operating near maximum power capture a selective advantage. However, treating power maximization as an unconstrained goal overlooked the stabilizing architecture of coupled transport networks: unconstrained dissipation leads to runaway depletion of essential reserves. What early maximum power formulations lacked was the reciprocal constraint provided by Onsager reciprocity and Le Chatelier-Braun moderation. Incorporating Onsager’s relations demonstrates that ecosystems do not merely maximize throughput; rather, intrinsic primary self-damping and coupled secondary pathways adaptively balance, stabilize, and throttle metabolic flows against environmental fluctuations (Choi et al. 1999; Choi et al. 2001).

## 7 Conclusion

Thermodynamics provides a consistent physical basis for integrating ecological models that are otherwise treated as phenomenological. In many respects, ecology faces a problem analogous to early thermal physics before statistical mechanics connected macroscopic state variables to microscopic dynamics, though ecology has the advantage of drawing directly on Boltzmann, Gibbs, Onsager, and Prigogine.

Statistical-mechanical configuration counts and macroscopic caloric budgets operate on different domains and should not be conflated; open living systems sustain local organization (*order* → *order*, *disorder* → *order*) only by exporting entropy across their boundaries (*(local order* ↔ *local order)* → *(global disorder)*). Moving from ill-defined whole-system accounting to a local, intensive control volume (*V ^‡^*) removes domain-scaling artifacts and clarifies why demographic interaction asymmetry does not contradict Onsager reciprocity. This is because demographic interaction coefficients (**A**) are not equivalent to physical conductivities, and at the statistical-ensemble scale of recurrent matter and energy cycling, dissipative coupling can be shown to be governed by symmetric, positive-definite conductivities (**L** ≻ 0) and specific entropy production (*σ* ≥ 0). Within this coupled transport framework, network cross-coupling moderates perturbations through direct self-damping (*L_ii_ >* 0) and cross-pathway redistribution (0 *< L_ii,_*_eff_ ≤ *L_ii_*), organizing communities along a biomass-capacitance versus turnover trade-off without invoking teleology. The companion paper (Choi 2026) explores the mapping between **A** and **L**, theoretically and empirically.

## Footnotes

1 This review is written for ecologists seeking a coherent synthesis of non-equilibrium thermodynamics, specifically in the Onsager-Prigogine tradition, with quantitative community ecology, where disparate terminologies and disciplinary conventions have historically obscured shared physical structure.

2 Landsberg’s (1961, 1972) representation of the laws of thermodynamics is amongst the most concise and clear as they are simple postulates of existence: - 0^th^ law: empirical temperature exists. - 1^st^ law: internal energy exists. Energy is conserved; it is not created nor destroyed, only transformed. - 2^nd^ law: *entropy* and absolute temperature exist. Every irreversible energy transformation generates *entropy*(Δ*S >* 0), degrading the free energy available to perform useful work. - 3^rd^ law: systems cannot attain a temperature of 0 Kelvin. - 4^th^ law: *extensive* (aggregate) and *intensive* (density) variables exist, for equilibrium states and at least some subset of near-equilibrium/non-equilibrium states.

3 In the linear near-equilibrium regime, Prigogine’s minimum entropy production theorem shows that unconstrained internal forces relax to minimize *σ*(**X**) = **X**^T^**LX** under fixed boundary forces (yielding vanishing internal fluxes), whereas Onsager’s (1931a, 1931b) and Onsager and Machlup’s (1953) principle of least dissipation of energy shows that realized fluxes at fixed forces minimize the Rayleighian 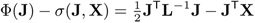, or equivalently maximize *σ −* Φ (Shiner et al. 1994; Martyushev et al. 2006; Verhás 2014). This paper focuses upon the linear range of applicability of Onsager’s relations, with appropriate caveats regarding far-from-equilibrium extensions.

4 The Le Chatelier-Braun principle, formulated between 1884 and 1888 (Gündüz et al. 2014), describes the empirical observation that physical-chemical systems tend to adjust their internal state when subjected to external perturbations so as to moderate or resist the imposed displacement. In coupled transport networks, this operates through two coupled responses combining direct primary self-damping (*L_ii_ >* 0) with secondary network buffering (0 *< L_ii,_*_eff_ *≤ L_ii_*). This general thermodynamic property in coupled irreversible networks is designated *Conjugate effects* to emphasize the constitutive mechanism provided by Onsager reciprocity, even in linear approximations.

5 The 4^th^ law, attributable to Landsberg (1972), is not universally acknowledged as a thermodynamic “law”. Though not controversial, it is important in the Onsager-Prigogine-Landsberg school and merits emphasis as it is not always appreciated or understood in the ecological literature. To emphasize this distinction, the qualification (*local, intensive*) is maintained throughout this work. When system volume is assumed constant, *intensive* and *extensive* quantities are frequently conflated, particularly in atmospheric applications. In ecological systems, macroscopic (*extensive*) biovolume variations of several orders of magnitude complicate naive applications of whole-system Onsager relations. Identifying standing biomass (or biovolume) as the active capacitive medium within local control volumes is a key to understanding these relations.

## References

Aoki, I. 1989. “Holological study of lakes from an entropy viewpoint: Lake Mendota.” Ecological Modelling 45 (2): 81–93.

Aoki, I. 1993. “Inclusive Kullback index: a macroscopic measure in ecological systems.” Ecological Modelling 66:289–299.

Aoki, I. 1995. “Entropy production in living systems: from organisms to ecosystems.” Thermochimica Acta 250:359–370.

Babloyantz, A. 1986. Molecules, Dynamics and Life: An Introduction to Self-Organization of Matter. Toronto: Wiley-Interscience.

Bell, G. 1982. The Masterpiece of Nature: The Evolution and Genetics of Sexuality. Berkeley: University of California Press.

Bertalanffy, L. von. 1950. “The theory of open systems in physics and biology.” Science (New York, N.Y.) 111:23–29.

Boltzmann, L. 1877. “Uber die Beziehung zwischen dem zweiten Hauptsatze der mechanischen Warmetheorie und der Wahrscheinlichkeitsrechnung respektive den Satzen uber das Warmegleichgewicht.” Wiener Berichte 76:373–435.

Briedis, D., and R. C Seagrave. 1984. “Energy dissipation in oscillating chemical reactions.” The Journal of Chemical Physics 80 (7): 3209–3217.

Byers, R. E., and R. I. C. Hansell. 1996. “Implications of semi-stable attractors for ecological modelling.” Ecological Modelling 89:59–65.

Carnot, S. 1824. Reflexions sur la Puissance Motrice du Feu et sur les Machines Propres a Developper Cette Puissance. Paris: Bachelier.

Chakrabarti, C. G., and K. Ghosh. 2009. “Non-equilibrium thermodynamics of ecosystems: Entropic analysis of stability and diversity.” Ecological Modelling 220 (16): 1950–1956.

Chapman, E. J., D. L. Childers, and J. J Vallino. 2016. “How the Second Law of Thermodynamics Has Informed Ecosystem Ecology through Its History.” BioScience 66:27–39.

Choi, J. S. 2002. “Dealing with uncertainty in a complicated thermodynamic world.” PhD diss., Doctoral dissertation.

Choi, J. S. 2019. “Thermodynamics in Ecology.”

Choi, J. S. 2022. “Reconstructing the decline of Atlantic Cod with the help of environmental variability in the Scotian Shelf of Canada” [in en]. *In Prep* (May): 2022.05.05.490753. 10.1101/2022.05.05.490753. https://www.biorxiv.org/content/10.1101/2022.05.05.490753v3.

Choi, J. S. 2026. Conjugates in Ecology. bioRxiv preprint.

Choi, J. S., Kenneth T Frank, William C Leggett, and Ken Drinkwater. 2004. “Transition to an alternate state in a continental shelf ecosystem.” Canadian Journal of Fisheries and Aquatic Sciences 61, no. 4 (April): 505–510. issn: 0706-652X. 10.1139/f04-079.

Choi, J. S., A. Mazumder, and R. I. C. Hansell. 1999. “Measuring perturbation in a complicated, thermodynamic world.” Ecological Modelling 117:143–158.

Choi, J. S., and B. C. Patten. 2001. “Sustainable development: lessons from the paradox of enrichment.” Ecosystem Health 7:163–178.

Clausius, R. 1850. “Ueber die bewegende Kraft der Warme und die Gesetze, welche sich daraus fur die Warmelehre selbst ableiten lassen.” Annalen der Physik 155 (3): 368–397.

Crooks, Gavin E. 1999. “Entropy production fluctuation theorem and the nonequilibrium work relation for free energy differences.” Physical Review E 60:2721–2726. 10.1103/PhysRevE.60.2721.

Drake, J. W., B. Charlesworth, D. Charlesworth, and J. F Crow. 1998. “Rates of spontaneous mutation.” Genetics 148 (4): 1667–1686.

Eigen, M., and P Schuster. 1979. The Hypercycle: A Principle of Natural Self-Organization. Berlin: Springer-Verlag.

Elith, J., S. J. Phillips, T. Hastie, M. Dudík, Y. E. Chee, and C. J Yates. 2011. “A statistical explanation of MaxEnt for ecologists.” Diversity and Distributions 17 (1): 43–57.

Elton, C. 1927. Animal Ecology. London: Sidgwick / Jackson.

Eppley, R. W., and B. J Peterson. 1979. “Particulate organic matter flux and planktonic new production in the deep ocean.” Nature 282 (5740): 677–680.

Fath, B. D., S. E. Jørgensen, B. C. Patten, and M Straškraba. 2004. “Ecosystem growth and development.” Biosystems 77:213–228.

Gibbs, J. W. 1902. Elementary Principles in Statistical Mechanics. New York: Charles Scribner’s Sons.

Gladyshev, G.P. 1978. “On the thermodynamics of biological evolution.” Journal of Theoretical Biology 75:425–441.

Glansdorff, P., G. Nicolis, and I Prigogine. 1974. “The thermodynamic stability theory of non-equilibrium states.” Proceedings of the National Academy of Sciences 71 (1): 197–199.

Glansdorff, P., and I. Prigogine. 1971. Thermodynamic Theory of Structure, Stability and Fluctuations. London: Wiley-Interscience.

Gould, S.J., and R.C Lewontin. 1979. “The spandrels of San Marco and the Panglossian paradigm: a critique of the adaptationist programme.” *Proceedings of the Royal Society of London*, Series B: Biological Sciences 205:581–598.

Grimm, V., and C Wissel. 1997. “Babel, or the ecological stability discussions: an inventory and analysis of terminology and a guide for avoiding confusion.” Oecologia 109:323–334.

Groot, S. R. de, and P Mazur. 1962. Non-Equilibrium Thermodynamics. Amsterdam: North-Holland Publishing Company.

Gündüz, Ö., and G Gündüz. 2014. “The phase space interpretation of the Le Chatelier-Braun principle and its generalization as a principle of natural philosophy.” Physics Essays 27 (3): 404–412. https://doi.org/10.

Halbach, U. 1973. “Life table data and population dynamics of the rotifer *Brachionus calyciflorus* Pallas as influenced by periodically oscillating temperature.”

Holdaway, R.J., A.D. Sparrow, and D.A Coomes. 2010. “Trends in entropy production during ecosystem development in the Amazon Basin.” Philosophical Transactions of the Royal Society of London B: Biological Sciences 365:1437–1447.

Hutchinson, G.E. 1959. “Homage to Santa Rosalia, or why are there so many kinds of animals? *The American Naturalist*, 93(870), pp.” 145–159.

Jaynes, E. T. 1957. “Information theory and statistical mechanics.” Physical Review 106:620–630. 10.1103/PhysRev.106.620.

Johnson, L. 1981. “The thermodynamic origin of ecosystems.” Canadian Journal of Fisheries and Aquatic Sciences 38 (5): 571–590.

Jørgensen, S. E. 2012. Integration of Ecosystem Theories: A Pattern. Dordrecht: 3rd edn. Springer.

Jørgensen, S. E., S. N. Nielsen, and H Mejer. 1995. “Emergy, environ, exergy and ecological modelling.” Ecological Modelling 77:99–109.

Katchalsky, A., and P. F Curran. 1965. Nonequilibrium Thermodynamics in Biophysics. Cambridge, MA: Harvard University Press.

Kay, J. J. 1991. “A nonequilibrium thermodynamic framework for discussing ecosystem integrity.” Environmental Management 15 (4): 483–495.

Kedem, O., and A Katchalsky. 1958. “Thermodynamic analysis of the permeability of biological membranes to non-electrolytes.” Biochimica et Biophysica Acta 27:229–246.

Kleiber, Max. 1947. “Body size and metabolic rate.” Physiological Reviews 27:511–541.

Kleidon, A. 2010. “A basic introduction to the thermodynamics of the Earth system far from equilibrium and maximum entropy production.” Philosophical Transactions of the Royal Society of London B: Biological Sciences 365:1303–1315.

Kolmogorov, Andrei N. 1941. “The local structure of turbulence in incompressible viscous fluid for very large Reynolds numbers.” Doklady Akademii Nauk SSSR 30:301–305.

Landsberg, P. T. 1961. Thermodynamics with Quantum Statistical Illustrations. New York: Interscience Publishers.

Landsberg, P. T. 1972. “The Fourth Law of Thermodynamics.” Nature 238:229–231.

Lindeman, R. L. 1942. “The trophic-dynamic aspect of ecology.” Ecology 23 (4): 399–417.

Lotka, A. J. 1922a. “Contribution to the energetics of evolution.” Proceedings of the National Academy of Sciences 8 (6): 147–151.

Lotka, A. J. 1922b. “Natural selection as a physical principle.” Proceedings of the National Academy of Sciences 8 (6): 151–154.

Lovelock, J. E. 1972. “Gaia as seen through the atmosphere.” Atmospheric Environment 6 (8): 579–580.

Lovelock, J. E., and L Margulis. 1974. “Atmospheric homeostasis by and for the biosphere: the Gaia hypothesis.” PhD diss.

Lurié, D., and J Wagensberg. 1979. “Non-equilibrium thermodynamics and biological growth and development.” Journal of Theoretical Biology 78:241–250.

MacArthur, R.H., and E.R Pianka. 1966. “On optimal use of a patchy environment.” The American Naturalist 100 (916): 603–609.

Mann, K.H. 1972. “Ecological energetics of the seaweed zone in a marine bay on the Atlantic Coast of Canada.” I. Zonation and biomass of seaweeds. *Marine Biology*, 12, pp.1–10.

Margalef, R. 1963. “On certain unifying principles in ecology.” The American Naturalist 97 (897): 357–374.

Margalef, R. 1968. Perspectives in Ecological Theory. Chicago: University of Chicago Press.

Martyushev, L. M., and V. D Seleznev. 2006. “Maximum entropy production principle in physics, chemistry and biology.” Physics Reports 426 (1): 1–45.

Matsuno, K. 1978. “Evolution of dissipative systems: a theoretical basis for Margalef’s principle on ecosystems.” Journal of Theoretical Biology 70:23–31.

Mayr, E. 1991. “The ideological resistance to Darwin’s theory of natural selection.” Proceedings of the American Philosophical Society 135:123–139.

Mazur, P. 1997. “Onsager’s reciprocal relations and thermodynamics of irreversible processes.” Periodica Polytechnica Chemical Engineering 41:197–204.

Meysman, Filip J. R., and Stijn Bruers. 2010. “Ecosystem functioning and maximum entropy production: a quantitative test of hypotheses.” Philosophical Transactions of the Royal Society of London B: Biological Sciences 365 (1545): 1405–1416.

Michaelian, Karo. 2005. “Thermodynamic stability of ecosystems.” Journal of Theoretical Biology 237 (3): 323–335.

Mikulecky, D. C. 1985. “Network thermodynamics in biology and ecology: an introduction.” In Ecosystem Theory for Biological Oceanography, edited by R. E. Ulanowicz and T. Platt, 213:163–175. Canadian Bulletin of Fisheries and Aquatic Sciences.

Miller, D. G. 1960. “Thermodynamics of irreversible processes.” The experimental verification of the Onsager reciprocal relations. *Chemical Reviews*, 60(1), pp. 15–37.

Müller, F. 1998. “Gradients in ecological systems.” Ecological Modelling 108:3–21.

Nicolis, G., and I Prigogine. 1977. Self-Organization in Nonequilibrium Systems: From Dissipative Structures to Order through Fluctuations. New York: John Wiley & Sons.

Nicolis, Gregoire, and Ilya Prigogine. 1989. Exploring Complexity: An Introduction. W. H. Freeman.

O’Neill, R.V., D.L. DeAngelis, J.B. Waide, and T.F.H Allen. 1986. A Hierarchical Concept of Ecosystems. Princeton, NJ. 253 p: Monographs in Population Biology 23. Princeton University Press.

Odum, E. P. 1969. “The strategy of ecosystem development.” Science 164 (3877): 262–270.

Odum, H. T. 1956. “Primary production in flowing waters.” Limnology and Oceanography 1 (2): 102–117.

Odum, H. T. 1996. Environmental Accounting: Emergy and Environmental Decision Making. New York: John Wiley & Sons.

Odum, H. T., and R. C. Pinkerton. 1955. “Time’s speed regulator: the optimum efficiency for maximum power output in physical and biological systems.” American Scientist 43 (2): 331–343.

Onsager, L., and S Machlup. 1953. “Fluctuations and irreversible processes.” Physical Review 91 (6): 1505–1512.

Onsager, Lars. 1931a. “Reciprocal Relations in Irreversible Processes. I.” Physical Review 37:405–426. 10.1103/PhysRev.37.405.

Onsager, Lars. 1931b. “Reciprocal Relations in Irreversible Processes. II.” Physical Review 38:2265–2279. 10.1103/PhysRev.38.2265.

Oster, George, Alan Perelson, and Aharon Katchalsky. 1971. “Network thermodynamics.” Nature 234:393–399.

Patten, Bernard C. 1978. “Systems approach to the concept of environment.” Ohio Journal of Science 78:206–222.

Peters, R. H. 1983. The Ecological Implications of Body Size. Cambridge: Cambridge University Press.

Peters, R. H. 1991. A Critique for Ecology. Cambridge: Cambridge University Press.

Planck, M. 1901. “Ueber das Gesetz der Energieverteilung im Normalspektrum.” Annalen der Physik 309 (3): 553–563.

Prigogine, I. 1967. Introduction to Thermodynamics of Irreversible Processes. New York: 3rd ed. Inter-science Publishers.

Prigogine, I., and R Defay. 1954. Chemical Thermodynamics. London: Longmans Green / Co.

Prigogine, I., G. Nicolis, and A Babloyantz. 1972. “Thermodynamics of evolution.” Physics Today 25 (11): 23–28.

Prigogine, Ilya. 1955. Thermodynamics of Irreversible Processes. New York: John Wiley / Sons.

Regier, H. A., and J. J Kay. 1996. “An heuristic model of transformations of the aquatic ecosystems of the Great Lakes-St.” Lawrence River Basin. *Journal of Aquatic Ecosystem Health*, 5(1), pp. 3–21.

Rook, G. A. W. 2009. “Review series on helminths, immune modulation and the hygiene hypothesis: the broader implications of the hygiene hypothesis.” PhD diss.

Russell, F. L., D. B. Zippin, and N. L Fowler. 2001. “Effects of white-tailed deer (*Odocoileus virginianus*) on plants, plant populations and communities: a review.” The American Midland Naturalist 146 (1): 1–26.

Schneider, E. D., and J. J Kay. 1994. “Life as a manifestation of the second law of thermodynamics.” Mathematical and Computer Modelling 19:25–48.

Schneider, E. D., and D Sagan. 2005. Into the Cool: Energy Flow, Thermodynamics, and Life. Chicago: University of Chicago Press.

Schrödinger, E. 1944. What is Life? The Physical Aspect of the Living Cell. Cambridge: Cambridge University Press.

Shiner, J. S., and S. Sieniutycz. 1994. “The chemical dynamics of biological systems: variational and extremal formulations.” Progress in Biophysics and Molecular Biology 62:203–221.

Smith, S. J., H. Bourdages, J. S. Choi, E. Dawe, J. S. Dunham, L. Gendron, D. Hardie, M. Moriyasu, D. Orr, and D. Roddick. 2012. “Technical guidelines for the provision of scientific advice on the precautionary approach for Canadian fish stocks: Section 7–invertebrate species.” DFO Can. Sci. Advis. Sec. Res. Doc 117.

Strachan, D. P. 1989. “Hay fever, hygiene, and household size.” BMJ 299 (6710): 1259–1260.

Strachan, D. P. 2000. “Family size, infection and atopy: the first decade of the ’hygiene hypothesis’.” PhD diss.

Tansley, A. G. 1935. “The use and abuse of vegetational terms and concepts.” Ecology 16 (3): 284–307.

Thomson, W. (Lord Kelvin). 1849. “An account of Carnot’s theory of the motive power of heat; with numerical results deduced from Regnault’s experiments on steam.” Transactions of the Royal Society of Edinburgh 16 (5): 541–574.

Turing, A. M. 1952. “The chemical basis of morphogenesis.” Philosophical Transactions of the Royal Society of London B: Biological Sciences 237 (641): 37–72.

Ulanowicz, R. E., S. E. Jørgensen, and B. D Fath. 2006. “Exergy, information and aggradation: an ecosystems reconciliation.” Ecological Modelling 198:520–524.

Ulanowicz, Robert E., and Bruce M. Hannon. 1987. “Life and the production of entropy.” Proceedings of the Royal Society of London B 232:181–192.

Van Valen, L. 1973. “A new evolutionary law.” Evolutionary Theory 1:1–30.

Van Valen, L. 1976. “Energy and evolution.” Evolutionary Theory 1:179–229.

Vanriel, P., and L Johnson. 1995. “Action principles as determinants of ecosystem structure: the autonomous lake as a reference system.” Ecology 76:1741–1757.

Verhás, J. 2014. “Gyarmati’s Variational Principle of Dissipative Processes.” Entropy 16 (4): 2362–2383. https://doi.org/10.

Washida, T. 1995. “Ecosystem configurations consequent on the maximum respiration hypothesis.” PhD diss.

Watson, Andrew J., and James E. Lovelock. 1983. “Biological homeostasis of the global environment: the parable of Daisyworld.” Tellus B 35:284–289.

Whittaker, R.H. 1953. “A consideration of climax theory: the climax as a population and pattern.” Ecological Monographs 23:41–78.

Wicken, J. S. 1980. “A thermodynamic theory of evolution.” Journal of Theoretical Biology 87 (1): 9–23.

Wiley, E. O., and D. R Brooks. 1982. “Victims of history: a nonequilibrium approach to evolution.” Systematic Zoology 31 (1): 1–24.

Yen, J. D. L., D. M. Paganin, J. R. Thomson, and R Mac Nally. 2014. “Thermodynamic extremization principles and their relevance to ecology.” Austral Ecology 39 (6): 619–632.

Yodzis, P. 1981. “The stability of real ecosystems.” Nature 289 (5799): 674–676.

Yodzis, P. 1988. “The indeterminacy of ecological interactions as perceived through perturbation experiments.” Ecology 69 (2): 508–515. https://doi.org/10.

Zaikin, A. N., and A. M Zhabotinsky. 1970. “Concentration wave propagation in two-dimensional liquid-phase self-oscillating system.” Nature 225 (5232): 535–537.

Zotin, A. I., and R. S Zotina. 1967. “Thermodynamic aspects of developmental biology.” Journal of Theoretical Biology 17 (1): 57–75.

